# Evaluation of targeted oxidative stress induced by oxygen-ozone gas mixtures *in vitro* after ischemic induction

**DOI:** 10.1101/2022.06.16.496274

**Authors:** Jessica Rodrigues Orlandin, Sarah Ingrid Pinto Santos, Luciana Cristina Machado, Paulo Fantinato Neto, Fabiana Fernandes Bressan, Naira Caroline Godoy Pieri, Kaiana Recchia, Meline de Paula Coutinho, Priscilla Avelino Ferreira Pinto, Annalisa Santucci, Valter Travagli, Carlos Eduardo Ambrosio

## Abstract

Encephalic vascular accident, or stroke, is the most common pathology of the central nervous system in humans. It is the second leading cause of death, as well as physical and cognitive disabilities, in developing countries. It is a vascular disorder that may present in an ischemic (more common) or hemorrhagic form. Ozone is a gas capable of oxidizing double bonds of organic molecules, thereby producing lipoperoxides and aldehydes. Stimulation of the immune and antioxidant system, and improvement in tissue vascularization and oxygenation, are few among several effects exerted by ozone. Ozone therapy has previously been shown to be effective in neuromodulation, neuroprotection, and nerve regeneration. The present study aimed to evaluate the effect of targeted mild ozone after inducing cerebral ischemia *in vitro*. The experiment was divided into two steps; in the first step, neuroblastoma lineage cells (SH-SY5Y) were subjected to 24 hours of hypoxia in an incubator culture chamber. Treatment with different concentrations of ozone (2–10 µg/mL), followed by an 2,5-diphenyl-2H-tetrazolium bromide assay, indicated a possible neuroregenerative effect at low concentrations. The same protocol was applied to canine amniotic membrane stem cells that were evaluated via colorimetric assay spectrophotometry, fluorescence microscopy, and flow cytometry. The metabolic conditions and cellular regeneration in cells at low ozone concentrations (3–8 µg/mL) correlated with lower levels of apoptosis and oxidative stress compared to cells not subjected to hypoxia. High concentrations of ozone (18–30 µg/mL) promoted an increase in rate of apoptosis and cell death. Therefore, we developed a novel protocol that mimics ozone therapy for ischemic stroke, using ozonized culture medium after hypoxia induction. Although more studies are needed to open new avenues for translational medicine, we conclude that ozone has a dose-dependent hormetic effect and can reverse the effect of ischemia *in vitro* at low concentrations.

## INTRODUCTION

A stroke is a neurological deficit caused by an altered brain circulation. This term includes injuries caused by hemodynamic and clotting disorders in arteries or veins (1) and is the most common pathology of the central nervous system. Recent studies indicate that stroke has become the second leading cause of death due to physical and cognitive disabilities in developed countries (2,3). According to a 2021 report from the American Heart Association, ischemic strokes account for 87% of stroke cases (4). The main risk factors for stroke are hypertension, obesity, sedentary lifestyle, stress, high cholesterol, and smoking (5).

In ischemic stroke, interrupting cellular oxidative metabolism reduces phosphate and glucose production, releasing neurotransmitters and increasing calcium and sodium levels. These factors lead to a reduction in neuronal metabolism and mitochondrial function; energy insufficiency; formation of arachidonic acid, prostaglandin, and leukotrienes; vasoconstriction; platelet aggregation; and poor microvasculature (6,7).

Ozone is a gas composed of three oxygen atoms, performing a specialized function of oxidizing double bonds of organic molecules. It is a powerful oxidant that immediately reacts with hydrophilic antioxidants in plasma, such as uric acid, ascorbic acid, and albumin, producing lipoperoxides and aldehydes. Ozone exhibits a hormetic effect; it can peroxidize the erythrocyte membrane and induce a change in its negative charge, causing minimal oxidative stress. Moreover, it improves circulation and oxygen delivery in ischemic tissues, enhances cell metabolism, modulates antioxidant enzymes, induces a slight activation of the immune system, and increases the release of growth factors. Ozone also induces platelet degranulation, stimulates the neuroendocrine system, activates neuroprotective strategies, protects against apoptosis induced via oxidative stress, and increases gene repair activity. When administered in adequate concentration, no side effects are observed (8–13).

Previous studies indicate that ozone therapy can be effective in neuromodulation, neuroprotection, and nerve regeneration (14–21). However, very few studies use the technique for stroke or other low cerebral perfusion syndromes (14,21,22).

It is worth noting that most studies use ozone as a preconditioning strategy before inducing hypoxia. This approach is not feasible since patients affected by this type of illness seek treatment only after the onset of symptoms and not as prevention. Our pioneering research efforts utilize ozone after hypoxia, with cells still in culture, without resuspending them for ozone treatment.

The present study aimed to evaluate the *in vitro* effects of mild oxidative stress caused by ozone after induction of cerebral ischemia, mimicking an ischemic stroke. Ozone therapy, when applied in adequate concentrations, improves oxygenation levels and cell metabolism, promoting cell regeneration. The experiment was conducted in two phases, using neuroblastoma cells and amniotic membrane stem cells.

Neuroblastoma lineage cells (SH-SY5Y) were cultured *in vitro*, subjected to hypoxia in an incubator culture chamber, and then treated with different ozone concentrations (2–30 µg/mL). As expected, 2,5-diphenyl-2H-tetrazolium bromide (MTT) assays showed a hormetic effect of ozone: low concentrations could promote cell regeneration, while high concentrations were harmful to cells.

Similar results were observed in the second step of our study, where canine amniotic membrane stem cells were subjected to the same hypoxia-ozonization protocol and evaluated by colorimetric assay spectrophotometry, fluorescence microscopy, and flow cytometry. These results indicated concentration-dependent effects and possibly neuro-regeneration at low concentrations.

Although more studies are needed, our results suggest that *in vitro*, low concentrations were able to reverse the damage caused by hypoxia.

Importantly, our study proposed an unprecedented protocol. In contrast to our study, most other studies use ozone as preconditioning. This is incompatible with what we see in clinical routine, wherein ozone is used as post-stroke therapy.

## MATERIALS AND METHODS

The experiment was divided into two phases performed with neuroblastoma lineage cells and with amniotic membrane stem cells, carried out in Brazil. This project was submitted to the Ethics Committee on the Use of Animals of the Faculty of Animal Science and Food Engineering of the University of São Paulo on August 3, 2018 and was approved through protocol CEUA FZEA-USP n. 3722230818 and CEUA IFSC-USP n. 8689190419.

For the first step, SH-SY5Y neuroblastoma lineage cells were grown in bottles in culture medium (DMEM-high glucose + 10% fetal bovine serum + 1% penicillin/streptomycin + 1% glutamine) and incubated at 37 ºC and 5% CO_2_, for about 3 or 4 days, until they reached about 90% of confluence. Cells were trypsinized and counted using Thoma Chamber, with trypan blue. Approximately 1×10^−4^ SH-SY5Y cells, diluted in 200 µL of culture medium, were plated per well in a 96-well culture plate. Four wells were also plated with the same number of cells on another plate, as a hypoxia control. The plates were incubated at 37 ºC and 5% CO_2_ for 24 hours. The experiment was carried out thrice in quadruplicates. After 24 hours, the culture plate was placed inside an incubation culture chamber. The completely sealed chamber was filled with about 10% CO_2_ + N_2_, and the cells were incubated for another 24 hours at 37 ºC. The control plate (no hypoxia group) was not subjected to the hypoxia protocol and remained incubated at 37 ºC and 5% CO_2_ for 24 hours. A difference in the color of the medium could be observed after 24 hours of hypoxia.

After 24 hours, 1.5 mL of culture medium was mixed with 1.5 mL of ozone in a 5 mL syringe and stirred for 5 minutes. Each syringe contained a different concentration of ozone (2, 5, 10, 15, 20, and 30 µg/mL); an additional syringe contained only O_2_, while no gas was added to another. The culture medium was removed from each well of the 96-well plate and replaced with ozonized culture medium containing different concentrations of ozone, according to the scheme below. The plate was incubated for another 24 hours. In the control plate, the culture medium was replaced with normal medium, with no addition of gas.

The colorimetric MTT assay was performed to verify cellular metabolic activity. After 24 hours, in a light-deprived environment, the ozonized medium was replaced with 200 µL of MTT (0.5 mg/mL diluted in PBS) per well. After a 90-minute incubation, MTT was removed, and 200 µL of DMSO was added per well. The absorbance was measured at 690 nm using a spectrophotometer (Multiskan Go Spectrophotometer - Thermo Fisher Scientific). The spectrophotometry results were transformed into a percentage, wherein the group not subjected to hypoxia represented 100%.

The experimental design is depicted in Figure 1.

**Figure 1.**
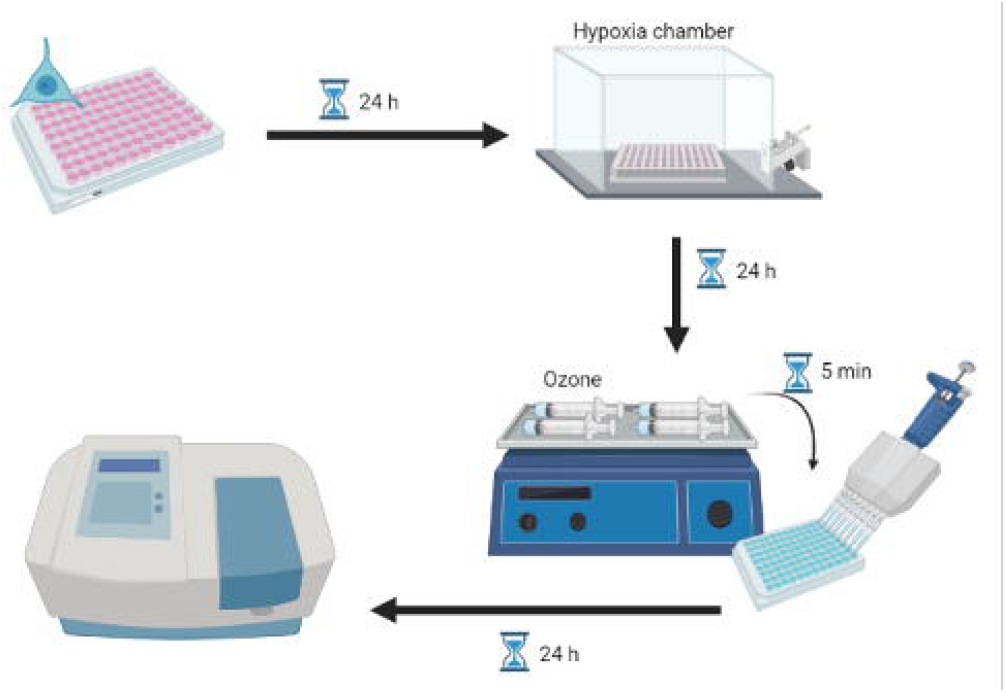
Experimental design of protocol for neuroblastoma cells. Cells were cultured *in vitro* and subjected to hypoxia. After 24 hours, the culture medium was replaced by ozonized medium at different concentrations for 5 minutes, incubated for another 24 hours, and then evaluated.

After satisfactory *in vitro* results with neuroblastoma cells, we decided to perform the same hypoxia-ozonation protocol with canine amniotic membrane stem cells, the same cells used in a previously published study (24). These mesenchymal stem cells are ease to obtain and can be discarded. They are easy to manipulate, and their proliferation is straightforward. These cells play an essential role in regenerative medicine, and having been previously characterized by our study group, prove to be safe for the application (25–28). After elective neutering surgery, the uterus of a bitch was sent to the laboratory at an early gestational stage. The amniotic membranes of eight fetuses were collected, as seen in Figure 2.

**Figure 2.**
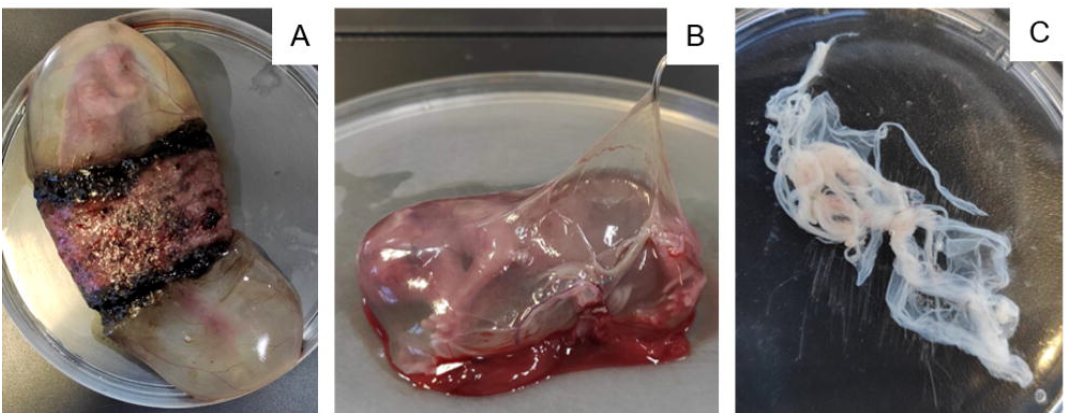
Isolation of canine amniotic membrane stem cells. A: Fetus in the first-third of the gestational period surrounded by the placenta. B: Dissection of amniotic membrane. C: Amniotic membrane in a culture plate.

Following the protocol by Orlandin et al. (24), membranes were washed in PBS with penicillin + streptomycin and incubated in 1% collagenase for 2 hours. After this period, they were centrifuged at 1200 rpm for 10 minutes. The pellet was resuspended in DMEM-high glucose + 10% fetal bovine serum + 1% penicillin/streptomycin + 0.5% glutamine, and the cell culture bottles were incubated at 37 °C and 5% CO_2_ for 3 days when they reached approximately 80% confluence.

For cell trypsinization, the medium was discarded and the bottle was washed with PBS. Tryple Express (3 mL; Gibco) were added. The bottle remained in an incubator for 2 minutes, to allow the cells to detach from the plate. Tryple Express was collected in a falcon tube, and the bottle was washed with the culture medium to help detach the remaining cells. This culture medium was collected in the same falcon and centrifuged at 1500 rpm for 5 minutes. Cells were picked and plated for an additional 4 days until they reached approximately 90% confluence. The trypsinization process was repeated, and the cells were counted using Thoma Chamber, with trypan blue, and harvested according to the specified assay.

For each assay, cells were harvested at different concentrations, using different plates, and at different volumes. Corresponding specifications are presented for each assay. Regardless, all of them were subjected to the same hypoxia protocol, as described below. All experiments were performed in triplicate and repeated three times. In addition to the experimental wells, separate wells were set aside for negative and positive control, for each protocol.

After 24 hours, the culture plate was placed inside a completely sealed incubator culture chamber, filled with about 10% CO_2_ + 90% N_2_, and incubated for another 24 hours. The control plate did not go through the hypoxia protocol and was incubated at 37 ºC and 5% CO_2_ for another 24 hours.

After 24 hours, a 20-mL syringe filled with 10 mL of culture medium was attached to an ozone generator (Ozone Generator O&L 3.0 RM – Ozone & Life Ltda – Brazil – ref 301140366), which injected another 10 mL of the gas into the syringe. They were stirred for 5 minutes. Each syringe contained a different concentration of ozone: 3, 8, 12, 18, and 30 µg/mL; additionally, a syringe with 10 mL of O_2_ and another without the addition of any gas (no gas group) were included.

The culture medium in each well was replaced by the ozonized medium at different concentrations. The plate was incubated for another 24 hours. In the control plate (no hypoxia group), the culture medium was replaced by a normal medium with no gas addition.

For the MTT assay, the above protocol was used. The absorbance was measured at 690 nm using a spectrophotometer (BioTek Epoch 2 microplate Spectrophotometer). The spectrophotometry results were transformed into percentage values, where the group not subjected to hypoxia represented 100%.

For the superoxide dismutase (SOD) and catalase (CAT) quantification, 1×10^−5^ cells were inoculated in a 6-well plate in 3 mL of culture medium per well in triplicate and subjected to the hypoxia-ozonation protocol, as mentioned above. The same plate was used for evaluating SOD (Superoxide Dismutase Colorimetric Activity Kit – Thermo Fisher Scientific – ref EIASODC) and CAT (Catalase Colorimetric Activity Kit –Thermo Fisher Scientific – ref EIACATC) activities, as only the medium was used to perform the SOD assay. In contrast, the cells were used to perform the CAT assay. The substrates were added, and the plate was incubated according to the manufacturer’s instructions. The absorbance was read at 450 nm and 520 nm by a spectrophotometer for SOD and CAT quantification, respectively.

For the apoptosis detection, we performed the TUNEL test, where 1×10^−3^ cells were harvested in a 96-well plate in 200 µL of culture medium per well in duplicate and subjected to the hypoxia-ozonation protocol, as mentioned above. Using the commercial Click-iT TUNEL (In Situ Cell Death Detection Kit, Fluorescein – Sigma-Aldrich – ref 11684795910) colorimetric detection kit, following the manufacturer’s instructions, the culture medium was removed and the cells were fixed with 4% paraformaldehyde and permeabilized with 0.1% Triton X-100 in 0.1% sodium citrate. TUNEL mixing solution (50 µL) was added to each sample and the positive and negative controls. The plate was incubated in an oven for 60 minutes, washed with PBS three times, and 200 µL of PBS was added for microscopic analysis. The cells were analyzed using a digital inverted EVOS microscope (Microscope EVOS™ FLc M5000 Imaging System – Thermo Fisher Scientific – ref AMEP4650) with a GFP laser (470 nm excitation/525 nm emission). The images were analyzed via fluorescence intensity quantification using ImageJ software, with intensity assignment between 0 and 200 per pixel.

The Image-iT® Lipid Peroxidation Kit has a lipid peroxidation sensitive agent. After oxidation, the fluorescence changes from red to green, allowing its quantification. For the lipid peroxidation assay, 2.5×10^−4^ cells were harvested in a 24-well plate in 1 mL of culture medium per well in triplicate and subjected to the hypoxia-ozonation protocol, as mentioned above. Image-iT® Lipid Peroxidation Sensor (10 µM; Image-iT Lipid Peroxidation Kit – Thermo Fisher Scientific – ref C10445) was added, and the plate was incubated at 37 ºC for 30 minutes in the dark. The culture medium was removed, and the plate was washed 3 times with PBS; 1 mL of PBS was added per well. As observable in Figure 15, the cells were analyzed with a digital inverted EVOS microscope, with a Texas Red laser (585 nm excitation/624 nm emission) and GFP (470 nm excitation/525 nm emission). The excitation and emission were adjusted for each probe. Cells were trypsinized and immediately read using a BD Accuri C6 flow cytometer (BD Accuri™ C6 Flow Cytometer – BD Biosciences – ref 7820018).

CellROX® Green is a fluorogenic probe that after oxidation, binds to DNA, rendering the nucleus and mitochondria fluorescent. For the ROS assay, 2.5×10^−4^ cells were harvested in a 24-well plate in 1 mL of culture medium per well in triplicate and subjected to the hypoxia-ozonation protocol, as mentioned above. CellROX Green (10 µM; CellROX Green Reagent, for oxidative stress detection – Thermo Fisher Scientific – ref C10444) reagent working dye solution was added to the culture medium of each well, followed by incubation at 37 ºC for 30 minutes in the dark. The culture medium was removed, and the plate was washed thrice with PBS; 1 mL of PBS was added per well. The cells were analyzed using a digital inverted EVOS microscope with a GFP laser (470 nm excitation/525 nm emission). Cells were trypsinized and read using a BD Accuri flow cytometer.

CellTracker™ Blue fluorescent probes cross the cell membrane and are transferred to daughter cells but not adjacent cells in a population. Therefore, they are an important tool for monitoring chemotaxis and cell proliferation. As it presents a reaction mediated by glutathione S-transferase, CellTracker Blue probes have also been used for GSH quantification (29,30). For the GSH assay, 1×10^−4^ cells were harvested in a 24-well plate in 1 mL of culture medium per well in duplicate and subjected to the hypoxia-ozonation protocol, as mentioned above. The culture medium was removed, and a solution containing 10 µM of CellTracker Blue (CellTracker Blue CMF2HC (Molecular Probes) – Thermo Fisher Scientific – ref C12881) in DMSO was added to each well. The plate was incubated for 30 minutes at 37 ºC in the dark. The solution was removed, and 1 mL of PBS was added per well. The cells were analyzed using a digital inverted EVOS microscope with a tgBFP laser (390 nm excitation/447 nm emission). Cells were trypsinized and read using a BD FACSARIA flow cytometer (BD FACSAria™ II Flow Cytometer Cell Sorter).

For all of the tests, data were analyzed using Prism 5.0. Tests for normality of residuals and homogeneity of variances were performed for each variable (body weight, clinical evaluation, and cerebral infarction volume). Data that did not meet the ANOVA assumptions were transformed. Statistical analysis of total infarct volume data was performed using the one-Way ANOVA test, a non-parametric statistic for purchasing groups, followed by Tukey’s Post-hoc test, considering the significance level (p<0.05). The results are expressed as mean ± standard error of the mean.

## RESULTS

### Spectrophotometry suggests cell regeneration at low ozone concentrations and apoptosis at high ozone concentrations

As expected, the neuroblastoma cells that suffered hypoxia, but received the culture medium without the addition of any gas, presented lower metabolic activity with respect to the control cells, which were not subjected to hypoxia. These results indicate that the protocol resulted in partial cell death, although it was not statistically significant (p>0.05). Cells treated with low concentrations of ozone (2 µg/mL) showed higher rates of cell viability than those that were subjected to hypoxia but did not receive any treatment or only received oxygen (p<0.05). This indicates that the treatment not only reversed the effects of ischemia but also promoted cell regeneration in these cells. Furthermore, cells treated with 2 µg/mL ozone were significantly similar to those not subjected to hypoxia and those treated with 5 and 10 µg/mL ozone (p>0.05), showing that these low concentrations are safe and maintain high rates of cell viability. Cells subjected to hypoxia that received no gas or only oxygen had the same cellular viability rate as those treated with 15 and 20 µg/mL of ozone (p>0.05). Finally, cells treated with 30 µg/mL of ozone showed a decrease in cell viability (p<0.05), indicating that high concentrations of ozone can harm cells.

Amniotic membrane stem cells (AMSC) treated with 3 µg/mL of ozone showed no significant difference compared to cells that did not undergo the hypoxia protocol (p>0.05). Additionally, these cells are similar to those that received no gas, received only oxygen, or were treated with 8 µg/mL ozone (p>0.05). Cells treated with low concentrations of ozone (2–8 µg/mL) showed better cell metabolism, compared to cell metabolism noted for other ozone treatments (12–30 µg/mL) (p<0.05), implying that these concentrations are safe and do not promote cell damage. Treating cells with 18 and 30 µg/mL of ozone harmed them, promoting cellular apoptosis compared to the cells in other groups (p<0,05). These results corroborate the results of the experiments conducted on our neuroblastoma cell lineages.

SOD analyses were performed on the culture medium and not on the cells. They were intended to mimic a serological quantification of SOD emitted by cells. SOD activity is associated with the cellular ability to scavenge free radicals. All ozone-treated groups had SOD levels comparable to cells not subjected to hypoxia (p>0.05). Cells that were subjected to hypoxia but received no gas or only oxygen had higher levels of SOD when compared to the group without hypoxia (p<0.05). It is important to emphasize that the commercial kit allows the quantification of SOD-1 and SOD-2, which are intracellular and mitochondrial enzymes, and SOD-3, which is extracellular and can be detected in serum, plasma, ascites, and synovial fluids. We tried to standardize a protocol for quantifying SOD in the culture medium to mimic the serological levels of the enzyme. Nonetheless, the results are inconsistent, and we do not consider them to be representative.

We observed a significant decrease in catalase rate following the increase in ozone concentration. Catalase levels of cells not subjected to hypoxia were significantly similar to those treated with no gas, only oxygen, and 3 and 8 µg/mL ozone (p>0.05). Cells treated with 30 µg/mL of ozone had the lowest levels of the intracellular enzyme, followed by cells of the groups treated with 18 and 12 µg/mL ozone, all of which presented significant differences between each other (p<0.05). This probably occurred due to the cytotoxic effect of ozone at high concentrations, as one of its by-products is hydrogen peroxide, a substance degraded by the peroxidase under study.

The spectrophotometry results have been presented in Figure 3.

**Figure 3.**
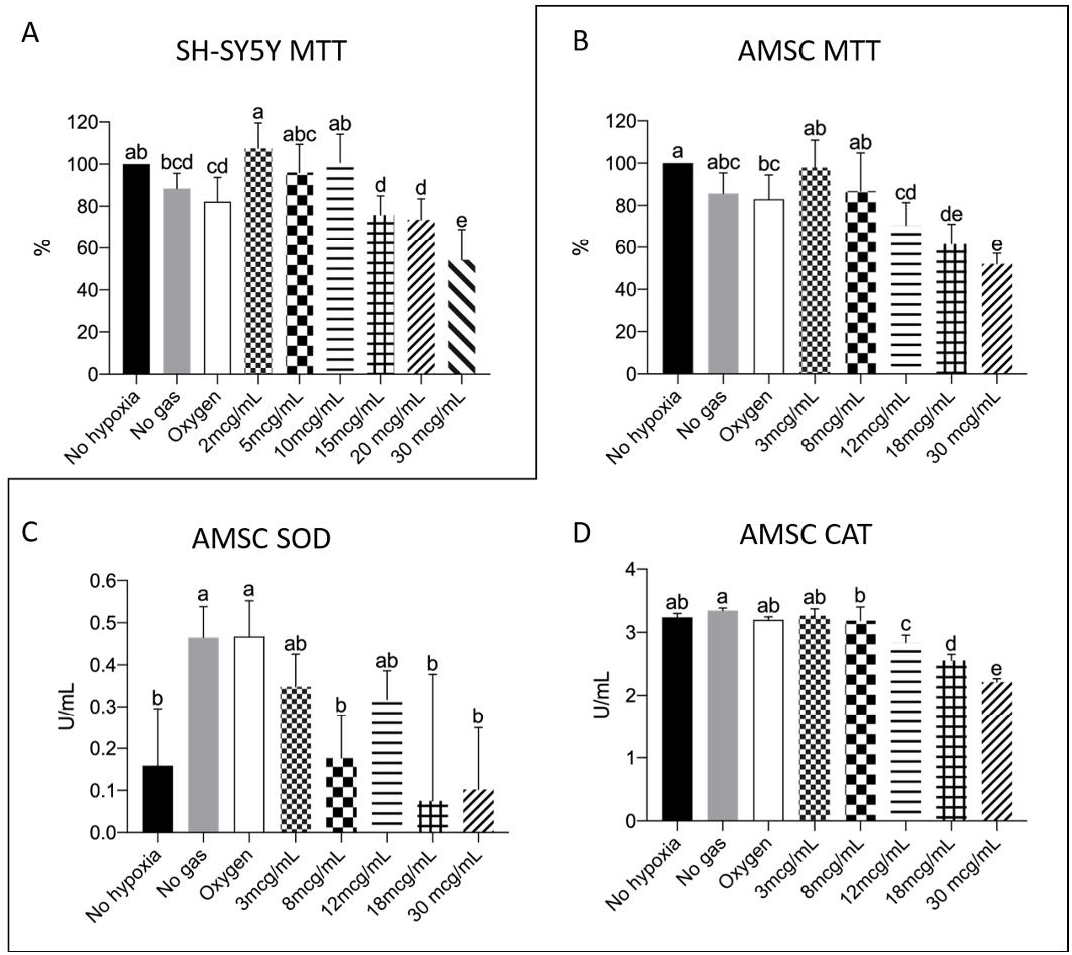
SH-SY5Y MTT (A) and AMSC MTT (A), SOD (B), and CAT (C) quantification, where p<0.05. During the MTT test, cells not subjected to the hypoxia protocol were considered 100%. We observed a pattern of reversal of the hypoxia effect and the promotion of cell regeneration at low ozone concentrations. Cellular apoptosis at high ozone concentrations was prominent in both kinds of cells (A and B). We also observed that SOD spectrophotometry results were inconsistent (C) and that high ozone concentrations promoted a decrease in catalase rate after hypoxia induction (D).

### Apoptosis detection assay corroborated the MTT data for high ozone concentrations promoting cell death

Due to material supply problems during the pandemic, we could not receive the TUNEL Kit in time to perform the tests. To this end, we borrowed two kits from a teacher at the institution. One of the kits was outdated and did not yield good images. Due to material limitations, cells grown in 96-microwell plates could not be subjected to flow cytometry. Therefore, the results could only be obtained from one sample from each group (Figure 4), and we could not perform duplicates or triplicates. Forty cells from each group were evaluated using ImageJ software for fluorescence rate (grey-value), resulting in the following graph.

**Figure 4.**
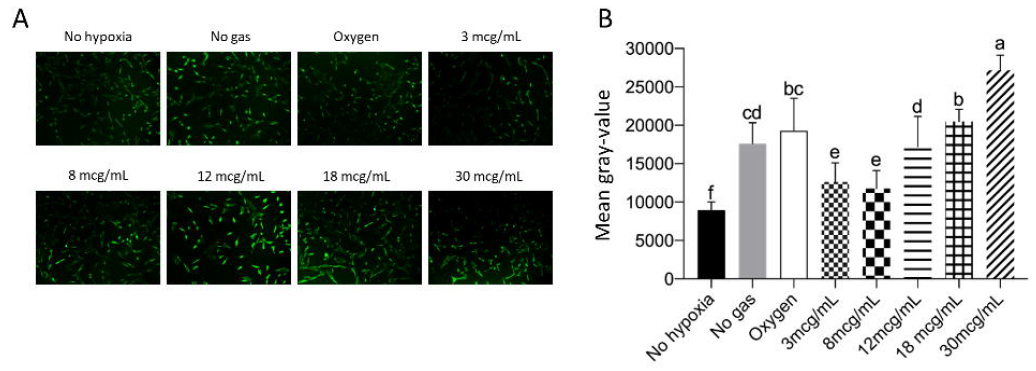
The higher the ozone level, the higher the rate of cell apoptosis. A: Representative pictures of cell apoptosis measured by TUNEL assay. C: Quantification of cell apoptosis (Gray-value, calculated by ImageJ), where p<0.05.

Cells that were not subjected to hypoxia showed the lowest levels of cellular apoptosis compared to the other groups (p<0.05). Cells treated with 3 or 8 µg/mL of ozone showed higher levels of cellular apoptosis than the cells in the no hypoxia group but lower levels than the cells in other groups (p<0,05). Cells subjected to hypoxia, but not receiving any gas, behaved similarly to those receiving oxygen (p>0.05). Nevertheless, they exhibited a lower rate of cellular apoptosis than cells of groups treated with 18 or 30 µg/mL (p<0.05). Cells treated with 30 µg/mL of ozone showed the highest rates of cell apoptosis compared to cells of all other groups (p<0.05).

These results are consistent with those of the MTT assay, wherein the mortality rate was higher in the groups treated with high ozone concentrations.

### No significant differences were observed in lipid peroxidation

The Image-iT® Lipid Peroxidation Kit has a lipid peroxidation sensitive agent. After oxidation, the fluorescence changes from red to green, allowing its quantification. As noted, the assessment of lipid peroxidation did not reveal significant differences between any of the groups (p>0.05). The results have been presented in Figure 5.

**Figure 5.**
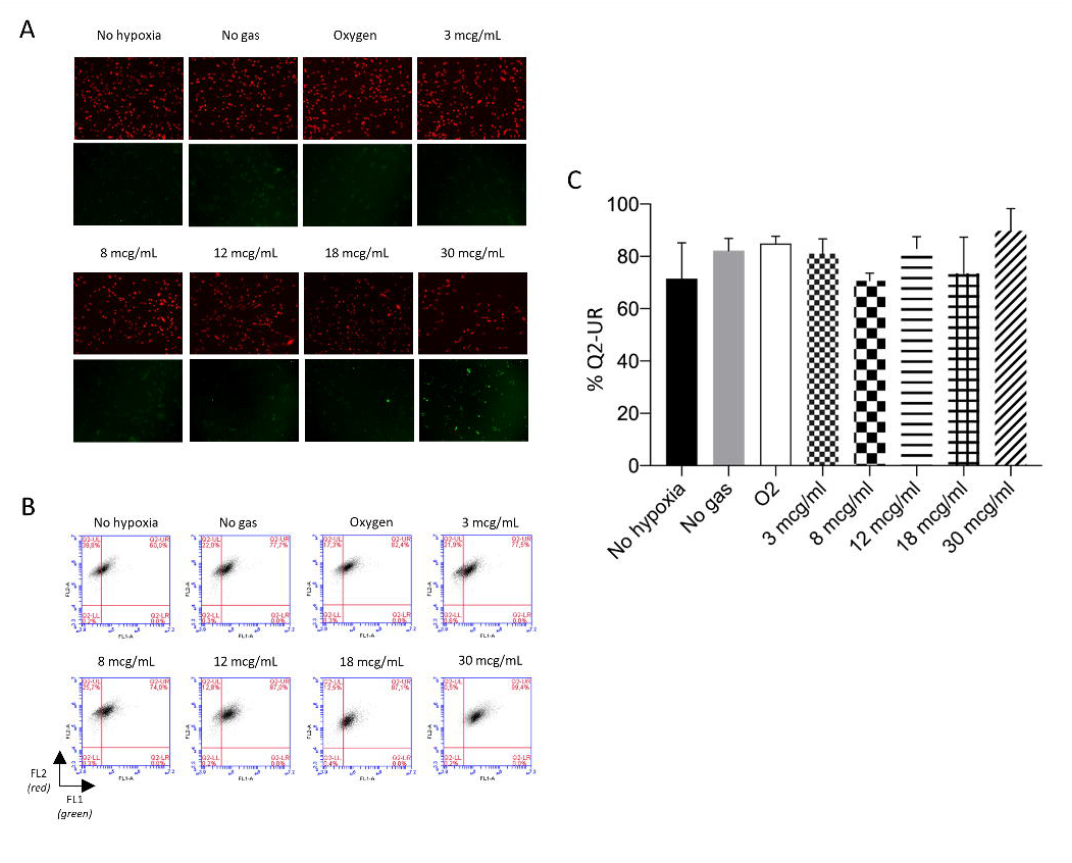
No treatment was of significant importance for lipid peroxidation. A: Representative pictures of lipidic peroxidation measured by Image-iT assay. B: Non-oxidized cells (red) and oxidized cells (green) measured by flow cytometry. C: Quantifying oxidized and non-oxidized cells (Q2-UR), where p<0.05.

### Low ozone concentration did not change ROS levels

CellROX® Green is a fluorogenic probe that after oxidation, binds to DNA, making the nucleus and mitochondria fluorescent. The results are presented in Figure 6.

**Figure 6.**
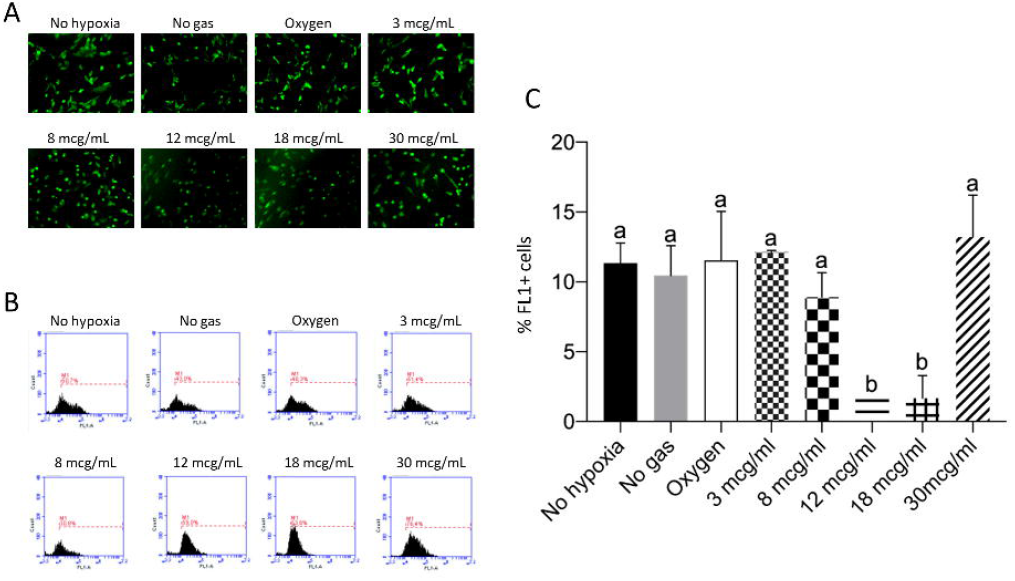
Low ozone concentrations do not increase the rate of reactive oxygen species (ROS). A: Representative pictures of ROS, as detected via CellROX Green assay. B: ROS examined via flow cytometry (FL1). C: Quantification of ROS levels, where p<0,05.

According to the analyses, low concentrations of ozone (3 and 8 µg/mL) did not show changes in ROS levels, compared to groups that were not subjected to hypoxia without gas and oxygen (p>0,05). Only the groups treated with 12 or 18 µg/mL of ozone presented reduced levels of ROS, compared to the other groups (p<0.05). ROS levels increased again in cells treated with 30 µg/mL ozone, equaling the levels noted for the other groups (p>0.05).

### Reduced Glutathione (GSH) levels remain unchanged across groups

The results presented below (Figure 7) show no significant differences between samples (p>0.05) for reduced GSH.

**Figure 7.**
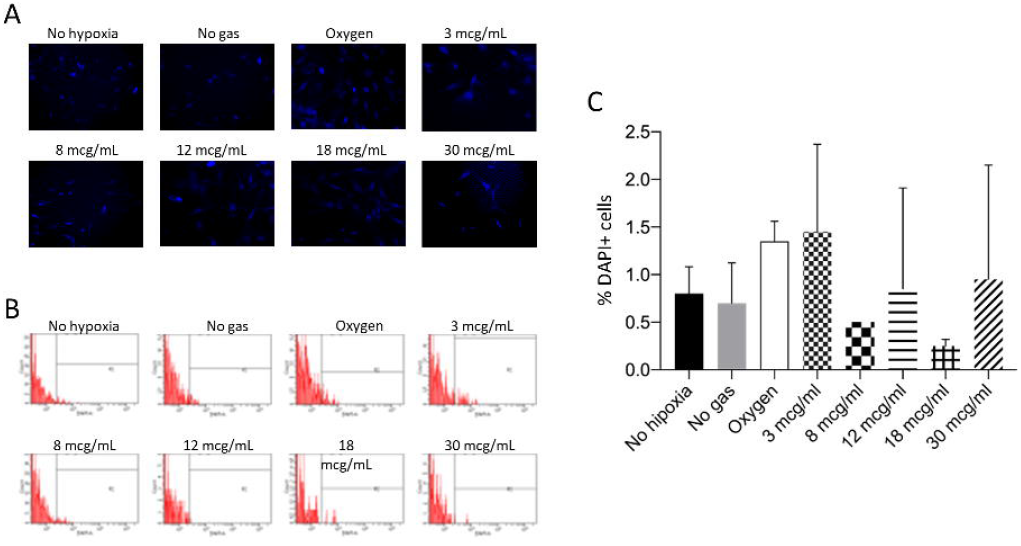
No treatment showed a significant difference in reduced glutathione level. A: Representative pictures of reduced glutathione (GSH) measured by CellTracker Blue assay. B: Reduced GSH measured by flow cytometry. C: Quantification of reduced GSH (DAPI), where p<0.05.

Importantly, these CellTracker™ Blue fluorescent probes are intended to stain daughter cells to screen the cell population. Few studies have used this probe to quantify GSH, owing to the reaction mediated by glutathione S-transferase. However, we reiterate that this is not the purpose recommended by the manufacturer (29,30).

Another critical point is that this technique was performed only in duplicate, with a very limited number of cells (1×10^3^ events). The data is unreliable and inconsistent. We do not recommend using the CellTracker™ Blue probe to quantify GSH in cells under the conditions discussed in this manuscript.

## DISCUSSION

In the present study, neuroblastoma cells and canine amniotic membrane stem cells were cultured *in vitro* and subjected to a 24-hour protocol in hypoxia to simulate an ischemic injury. Subsequently, the culture medium of cells was treated with different ozone concentrations, and the cellular response was evaluated.

Although ozone-mediated oxidative stress can contribute to the pathogenesis of neurodegenerative diseases (31), it is also known that mild, controlled, and targeted oxidative stress can be beneficial and can improve performance (32). Therefore, ozone is said to have a dose-dependent hormetic effect (10,12,33–38).

Corroborating the results of these studies, an MTT assay of neuroblastoma cells treated with post-hypoxia ozone indicated that low concentrations of ozone (2–10 µg/mL) were able to repair the effects of ischemia and promoted cell regeneration. This result is consistent with those of the MTT assay for canine amniotic membrane stem cells that underwent the same protocol. The group treated with 3–8 µg/mL showed improved cell metabolism. The results of TUNEL assay indicated that apoptosis rates are lower in cells treated with low ozone concentration (3–8 µg/mL). In all these trials, the harmful effects of ozone were observed at high concentrations (18–30 µg/mL), which promoted higher rates of apoptosis and cell death.

An increase in SOD levels after ozone treatment has been described previously (21,39,40). We attempted to develop a protocol for measuring SOD levels in a cell culture medium using the SOD Colorimetric Activity Kit (Thermo Fisher Scientific) to simulate serological levels of the enzyme. However, our results were inconsistent, and we could not conclusively comment on whether ozone-treated groups showed an increase or decrease in SOD levels. We recommend that such analysis be performed with these cells in the future, according to the manufacturer’s guidelines.

In contrast with some prior studies (21,39), we observed a progressive decrease in CAT levels as we increased the ozone concentration. CAT levels are an indicator of mitochondrial oxidative stress. Therefore, we expected increased CAT levels proportional to the increase in ozone concentration. Nevertheless, the same studies demonstrated a decrease in SOD and CAT levels after ischemia/reperfusion (IR). We reinforce here that our study involved usage of post-hypoxia ozone, and we believe that this may explain the observed inversion of CAT levels. We also suggest that the unexpected levels of CAT could be due to the reduction in the number of cells caused by gradual cell death with increasingly higher ozone doses.

As reported in other studies (41,42), we expected to observe an increase in lipid peroxidation corresponding to an increase in ozone concentration. Instead, in our study, the evaluation of lipid peroxidation (LPO) did not result in significant increase among any of the groups studied.

The quantification of ROS levels after hypoxia revealed no difference in the control and low ozone-treated groups (3 and 8 µg/mL). The groups treated with 12 or 18 µg/mL of ozone showed a significant decrease in ROS levels, which increased again in the group treated with 30 µg/mL, still being comparable with ROS levels noted for other groups. We expected to observe an increase in ROS levels, as it is one of the by-products of ozone, and the higher its concentration, the greater should be the production of ROS (43–47).

To quantify levels of reduced GSH, we used a protocol detailed in King et al. (29) and Casaril et al. (30), using CellTracker™ Blue fluorescent probes (Thermo Fisher Scientific). We restate that these probes, according to the manufacturer, are intended to stain daughter cells to screen a cell population. The data obtained were unreliable and inconsistent. We do not recommend using CellTracker™ Blue probe to quantify GSH in cells under the conditions discussed in this manuscript.

Few studies have evaluated the effect of ozone as a treatment for *in vitro* ischemia. Most studies used ozone for preconditioning cells, which were subjected to hypoxia only after treatment with the gas (39,40,48–53). Similar to our study, Cai et al. (21) used ozone after inducing hypoxia. However, they trypsinized the cells to perform the ozonation and re-plated them at a concentration stipulated for all groups.

In our study, all groups were initially plated at the same cell concentration. The culture plates were subjected to hypoxia. The cells themselves were not ozonized, but the cell culture medium was. We attempted to re-create the events of an ischemic stroke, wherein brain cells do not receive the gas, and instead, receive its by-products via the bloodstream. In addition, in real life, hardly any patient has sought treatment before disease onset. Patients who have suffered from ischemia and have survived seek therapies to minimize its sequelae. A literature survey did not reveal any publication using this protocol. Therefore, our study has pioneered the protocol for mimicking the *in vitro* consequences to a tissue treated with ozone therapy after suffering hypoxia.

Some professionals are against the practice of ozone therapy, owing to concerns regarding the toxicity of ozone at any concentration; however, such a perspective disregards the foundations of physiology and toxicology, as well as the advances in research and science. (46). Ozone therapy has been scientifically proven to be effective and safe, and thus, it should be considered a practical alternative by the relevant practitioners and researchers. Many of the studies presented on this topic have incomplete data, which impedes the standardization of the technique and the results obtained (46,54). We believe that these results should be shared and discussed to understand the technique and its mechanisms.

One of the recently raised hypotheses about the mechanisms of ozone therapy relates to its effect on blood cells and the extracellular matrix of the intravascular compartment (47). The interaction of ozone with blood cells and blood plasma during major autohemotherapy (when the blood of the ozonated patient is reinfused intravenously), resulting in LPO, primarily 4-hydroxynonenal (4-HNE); 4-HNE activates a cascade in the vascular endothelium, resulting in the release of nitric oxide, which acts as a vasodilator, and the upregulation of oxidative stress proteins and antioxidant enzymes to protect the vascular matrisome. With the intent of expanding such research efforts for application in clinical routine and considering that ischemic stroke is primarily a vascular disease, we believe that these factors should be examined in future research.

## CONCLUSION

To conclude, we reported a novel protocol, which involves using ozone after cell damage, as doing so is closer to the real-world scenario experienced by patients.

It is possible to perform *in vitro* post-hypoxia ozone therapy in cell cultures, without the need to resuspend them, simulating an ischemic stroke.

*In vitro*, low concentrations of ozone (2–10 µg/mL) enhanced the cellular metabolism of a neuronal cell line, which may be suggestive of post-hypoxia cell regeneration and reversal of cellular damage caused by hypoxia. High ozone concentrations (above 18 µg/mL) *in vitro* can cause cell damage.

Therefore, ozone exhibits a dose-dependent hormetic effect, and at low concentrations, is capable of reversing the effect of ischemia *in vitro*.

Although further studies are warranted, the use of ozone may be a promising therapeutic alternative for treating sequelae in stroke patients. Nevertheless, the concentration of ozone is an essential factor to be considered before prescribing treatment, especially in post-hypoxia conditions, wherein the cells are already damaged.

## ACKNOWLEDGMENTS

We would like to thank CAPES (PrInt No.88887.371111/2019-00) and FAPESP (No. 2018/24552-6) for their institutional support and Ozone & Life Ltda for providing us with the ozone generator used in our experiments. We also thanks Professor Fernando Fernandes Paiva and his team at Instituto de Física de São Carlos (IFSC-USP); the doctors Maria Serena Milella and Jasmine Ercoli; and the Professor Federica Pessina, Luana Ricci Paulesu e Maria Frosini from the University of Siena for their support.

## Notes

### Competing Interest Statement

The authors have declared no competing interest.

## REFERENCES

1. Klijn CJ, Kappelle LJ. Haemodynamic stroke: clinical features, prognosis, and management. The Lancet Neurology. 2010;9(10):1008–17. Available from: http://www.ncbi.nlm.nih.gov/pubmed/20864053

2. Jones F, Riazi A. Self-efficacy and self-management after stroke: a systematic review. Disability and Rehabilitation. 2011;33(10):797–810. Available from: http://www.ncbi.nlm.nih.gov/pubmed/20795919

3. Doyle PJ. Measuring health outcomes in stroke survivors. Archives of Physical Medicine and Rehabilitation. 2002;83:S39–43. Available from: http://linkinghub.elsevier.com/retrieve/pii/S0003999302006238

4. Virani SS, Alonso A, Aparicio HJ, Benjamin EJ, Bittencourt MS, Callaway CW, et al. Heart Disease and Stroke Statistics—2021 Update. Circulation. 2021;E254–743. Available from: https://www.ahajournals.org/doi/abs/10.1161/CIR.0000000000000950

5. Sumer M, Ozdemir I, Erturk O. Progression in acute ischemic stroke: Frequency, risk factors and prognosis. Journal of Clinical Neuroscience. 2003;10(2):177–80. Available from: https://www.sciencedirect.com/science/article/pii/S0967586802003259

6. Lim SH, Lee JS, Lee JI, Im S, Ko YJ, Kim HW. The quantitative assessment of functional impairment and its correlation to infarct volume in rats with transient middle cerebral artery occlusion. Brain Research. 2008;1230:303–9. Available from: http://www.ncbi.nlm.nih.gov/pubmed/18675259

7. Menzies SA, Hoff JT, Betz AL. Middle Cerebral Artery Occlusion in Rats. Neurosurgery. 1992;31(1):100–7. Available from: https://academic.oup.com/neurosurgery/article/31/1/100/2751985

8. Sagai M, Bocci V. Mechanisms of Action Involved in Ozone Therapy: Is healing induced via a mild oxidative stress? Medical Gas Research. 2011;1(1):29. Available from: http://www.ncbi.nlm.nih.gov/pubmed/22185664

9. Travagli V, Zanardi I, Silvietti A, Bocci V. A physicochemical investigation on the effects of ozone on blood. International Journal of Biological Macromolecules. 2007;41(5):504–11. Available from: https://www.sciencedirect.com/science/article/pii/S0141813007001675

10. Bocci V, Borrelli E, Travagli, V, Zanardi I. The ozone paradox: Ozone is a strong oxidant as well as a medical drug. Medicinal Research Reviews. 2009;29(4):646–82. Available from: http://www.ncbi.nlm.nih.gov/pubmed/19260079

11. Johnson JA, Johnson DA, Kraft AD, Calkins MJ, Jakel RJ, Vargas MR, et al. The Nrf2-ARE Pathway. Ann N Y Acad Sci. 2008;1147(1):61–9. Available from: http://www.ncbi.nlm.nih.gov/pubmed/19076431

12. Bocci V, Zanardi I, Travagli V. Oxygen/ozone as a medical gas mixture. A critical evaluation of the various methods clarifies positive and negative aspects. Medical Gas Research. 2011;1:1–6. Available from: /pmc/articles/PMC3231820/?report=abstract

13. Tufekci KU, Civi Bayin E, Genc S, Genc K. The Nrf2/ARE Pathway: A Promising Target to Counteract Mitochondrial Dysfunction in Parkinson’s Disease. Parkinson’s Disease. 2011;2011:1–14. Available from: http://www.ncbi.nlm.nih.gov/pubmed/21403858

14. Frosini M, Contartese A, Zanardi I, Travagli V, Bocci V. Selective ozone concentrations may reduce the ischemic damage after a stroke. Free Radical Research. 2012;46(5):612–8. Available from: http://www.ncbi.nlm.nih.gov/pubmed/22263539

15. Fuccio C, Luongo C, Capodanno P, Giordano C, Scafuro MA, Siniscalco D, et al. A single subcutaneous injection of ozone prevents allodynia and decreases the over-expression of pro-inflammatory caspases in the orbito-frontal cortex of neuropathic mice. Eur J Pharmacol. 2009;603(1–3):42–9. Available from: http://www.ncbi.nlm.nih.gov/pubmed/19100257

16. Ozbay I, Ital I, Kucur C, Akcılar R, Deger A, Aktas S, et al. Effects of ozone therapy on facial nerve regeneration. Braz J Otorhinolaryngol. 2017;83(2):168–75. Available from: http://www.bjorl.org

17. Ozturk O, Tezcan AH, Adali Y, Yildirim CH, Aksoy O, Yagmurdur H, et al. Effect of ozone and methylprednisolone treatment following crush type sciatic nerve injury. Acta Cirurgica Brasileira. 2016;31(11):730–5. Available from: http://www.scielo.br/scielo.php?script=sci_arttext&pid=S0102-86502016001100730&lng=en&tlng=en

18. Somay H, Emon ST, Uslu S, Orakdogen M, Meric ZC, Ince U, et al. The Histological Effects of Ozone Therapy on Sciatic Nerve Crush Injury in Rats. World Neurosurgery. 2017;105:702–8. Available from: https://www.sciencedirect.com/science/article/pii/S187887501730863X

19. Yucesoy T, Kutuk N, Canpolat DG, Alkan A. Comparison of Ozone and Photo-Biomodulation Therapies on Mental Nerve Injury in Rats. Journal of Oral and Maxillofacial Surgery. 2017;75(11):2323–32. Available from: http://www.ncbi.nlm.nih.gov/pubmed/28529151

20. Tricarico G, Isakovic J, Song M, Rustichelli F, Travagli V, Mitrecic D. Ozone influences migration and proliferation of neural stem cells in vitro. Neurosci Lett. 2020;739. Available from: https://pubmed.ncbi.nlm.nih.gov/32947004/

21. Cai H, Tao X, Zheng L, Huang L, Peng Y, Liao R, et al. Ozone alleviates ischemia/reperfusion injury by inhibiting mitochondrion-mediated apoptosis pathway in SH-SY5Y cells. Cell Biology International. 2020;44(4):975–84. Available from: https://onlinelibrary.wiley.com/doi/abs/10.1002/cbin.11294

22. Clavo B, Catalá L, Pérez JL, Rodríguez V, Robaina F. Ozone Therapy on Cerebral Blood Flow: A Preliminary Report. Evid Based Complement Alternat Med. 2004;1(3):315–9. Available from: http://www.ncbi.nlm.nih.gov/pubmed/15841265

23. Zhou N bao, Fu Z jian, Sun T. Effects of different concentrations of oxygen – ozone on rats’ astrocytes in vitro. Neuroscience Letters. 2008;441(2):178–82.

24. Orlandin JR, Gomes I da S, Leandro S de FS, Cagnim AF, Casals JB, Carregaro AB, et al. Treatment of Chronic Spinal Cord Injury in Dogs Using Amniotic Membrane-Derived Stem Cells: Preliminary Results. Stem Cells and Cloning: Advances and Applications. 2021;14:39–49. Available from: https://www.dovepress.com/treatment-of-chronic-spinal-cord-injury-in-dogs-using-amniotic-membran-peer-reviewed-fulltext-article-SCCAA

25. Cardoso M, Pinheiro A, Vidane A, Casals J, de Oliveira V, Gonçalves N, et al. Characterization of teratogenic potential and gene expression in canine and feline amniotic membrane-derived stem cells. Reproduction in Domestic Animals. 2017;52:58–64. Available from: http://doi.wiley.com/10.1111/rda.12832

26. Vidane A, Pinheiro A, Casals J, Passarelli D, Hage M, Bueno R, et al. Transplantation of amniotic membrane-derived multipotent cells ameliorates and delays the progression of chronic kidney disease in cats. Reproduction in Domestic Animals. 2017;52:316–26. Available from: http://www.ncbi.nlm.nih.gov/pubmed/27774657

27. Ambrosio C, Vidane A, Souza A, Sampaio R, Bressan F, Pieri N, et al. Cat amniotic membrane multipotent cells are nontumorigenic and are safe for use in cell transplantation. Stem Cells and Cloning: Advances and Applications. 2014;7:71. Available from: http://www.ncbi.nlm.nih.gov/pubmed/25249758

28. Pinheiro A de O, Lara VM, Souza AF, Casals JB, Bressan FF, Neto PF, et al. Characterization and Immunomodulation of Canine Amniotic Membrane Stem Cells. Stem Cells and Cloning: Advances and Applications. 2020;13:43–55. Available from: https://www.dovepress.com/characterization-and-immunomodulation-of-canine-amniotic-membrane-stem-peer-reviewed-fulltext-article-SCCAA

29. King N, Korolchuk S, McGivan JD, Suleiman MS. A new method of quantifying glutathione levels in freshly isolated single superfused rat cardiomyocytes. J Pharmacol Toxicol Methods. 2004;50(3):215–22. Available from: https://pubmed.ncbi.nlm.nih.gov/15519908/

30. Casaril AM, Segatto N, Simões L, Paschoal J, Domingues M, Vieira B, et al. Neuroprotective Effect of 3-[(4-Chlorophenyl)selanyl]-1-methyl-1H-indole on Hydrogen Peroxide-Induced Oxidative Stress in SH-SY5Y Cells. Neurochemical Research. 2021;46(3):535–49. Available from: https://link.springer.com/article/10.1007/s11064-020-03190-0

31. Moldogazieva NT, Mokhosoev IM, Mel’Nikova TI, Porozov YB, Terentiev AA. Oxidative Stress and Advanced Lipoxidation and Glycation End Products (ALEs and AGEs) in Aging and Age-Related Diseases. Oxidative Medicine and Cellular Longevity. 2019;2019. Available from: /pmc/articles/PMC6710759/

32. Berry R, López-Martínez G. A dose of experimental hormesis: When mild stress protects and improves animal performance. Comparative Biochemistry and Physiology Part A: Molecular & Integrative Physiology. 2020;242:110658.

33. Bocci VA, Zanardi I, Travagli V. Ozone acting on human blood yields a hormetic dose-response relationship. Journal of Translational Medicine. 2011;9(1):66. Available from: https://translational-medicine.biomedcentral.com/articles/10.1186/1479-5876-9-66

34. Bocci V. The Potential Toxicity of Ozone: Side Effects and Contraindications of Ozonetherapy. In: OZONE. Dordrecht: Springer Netherlands; 2010. p. 75–84. Available from: http://link.springer.com/10.1007/978-90-481-9234-2_7

35. Bocci V. How Does Ozone Act? How and Why Can We Avoid Ozone Toxicity? In: OZONE. Dordrecht: Springer Netherlands; 2010. p. 17–26. Available from: http://link.springer.com/10.1007/978-90-481-9234-2_4

36. Bocci V. Physical-Chemical Properties of Ozone – Natural Production of Ozone: The Toxicology of Ozone. In: OZONE. Dordrecht: Springer Netherlands; 2010. p. 1–4. Available from: http://link.springer.com/10.1007/978-90-481-9234-2_1

37. Bocci V. Is it true that ozone is always toxic? The end of a dogma. Toxicology and Applied Pharmacology. 2006;216(3):493–504. Available from: https://pubmed.ncbi.nlm.nih.gov/16890971/

38. Frosini M, Contartese A, Zanardi I, Travagli V, Bocci V. Selective ozone concentrations may reduce the ischemic damage after a stroke. Free Radical Research. 2012;46(5):612–8. Available from: http://www.tandfonline.com/doi/full/10.3109/10715762.2012.659247

39. Onal O, Yetisir F, Sarer AES, Zeybek ND, Onal CO, Yurekli B, et al. Prophylactic Ozone Administration Reduces Intestinal Mucosa Injury Induced by Intestinal Ischemia-Reperfusion in the Rat. Mediators of Inflammation. 2015;2015. Available from: /pmc/articles/PMC4487723/

40. RodriLuez ZZ, Guanche D, Álvarez RG, Rosales FH, Alonso Y, Schulz S. Preconditioning with ozone/oxygen mixture induces reversion of some indicators of oxidative stress and prevents organic damage in rats with fecal peritonitis. Inflamm Res. 2009;58(7):371–5. Available from: https://pubmed.ncbi.nlm.nih.gov/19274439/

41. Laszczyca P, Kawka-Serwecińska E, Witas I, Dolozych B, Falkus B, Mekail A, et al. Lipid peroxidation and activity of antioxidative enzymes in the rat model of ozone therapy. Materia Medica Polona. 1996;28(4):155–60. Available from: https://pubmed.ncbi.nlm.nih.gov/9308339/

42. Clavo B, Rodríguez-Esparragón F, Rodríguez-Abreu D, Martínez-Sánchez G, Llontop P, Aguiar-Bujanda D, et al. Modulation of Oxidative Stress by Ozone Therapy in the Prevention and Treatment of Chemotherapy-Induced Toxicity: Review and Prospects. Antioxidants. 2019;8(12):588. Available from: https://www.mdpi.com/2076-3921/8/12/588

43. Larini A, Bocci V. Effects of ozone on isolated peripheral blood mononuclear cells. Toxicology in Vitro. 2005;19(1):55–61.

44. Peralta C, Xaus C, Bartrons R, Leon OS, Gelpi E, Roselló-Catafau J. Effect of ozone treatment on reactive oxygen species and adenosine production during hepatic ischemia-reperfusion. Free Radic Res. 2000;33(5):595–605. Available from: https://pubmed.ncbi.nlm.nih.gov/11200091/

45. Bocci V, Valacchi G, Corradeschi F, Aldinucci C, Silvestri S, Paccaginini E, et al. Studies on the biological effects of ozone: 7. Generation of reactive oxygen species (ROS) after exposure of human blood to ozone. Journal of Biological Regulators & Homeostatic Agents. 1998;12(3):67–75. Available from: https://pubmed.ncbi.nlm.nih.gov/9795834/

46. Re L. Ozone in Medicine: A Few Points of Reflections. Frontiers in Physiology. 2022;13:103.

47. Tricarico G, Travagli V. The Relationship between Ozone and Human Blood in the Course of a Well-Controlled, Mild, and Transitory Oxidative Eustress. Antioxidants (Basel). 2021;10(12):1946–60. Available from: https://pubmed.ncbi.nlm.nih.gov/34943049/

48. Koca K, Yurttaş Y, Yildiz C, Çayci T, Uysal B, Korkmaz A. Effect of hyperbaric oxygen and ozone preconditioning on oxidative/nitrosative stress induced by tourniquet ischemia/reperfusion in rat skeletal muscle. Acta Orthop Traumatol Turc. 2010;44(6):476–83. Available from: https://pubmed.ncbi.nlm.nih.gov/21358255/

49. Ekici S, Doan Ekici AI, Öztürk G, Benli Aksungar F, Sinanolu O, Turan G, et al. Comparison of melatonin and ozone in the prevention of reperfusion injury following unilateral testicular torsion in rats. Urology. 2012;80(4):899–906. Available from: https://pubmed.ncbi.nlm.nih.gov/22950989/

50. Merin O, Attias E, Elstein D, Schwalb H, Bitran D, Zimran A, et al. Ozone administration reduces reperfusion injury in an isolated rat heart model. J Card Surg. 2007;22(4):339–42. Available from: https://pubmed.ncbi.nlm.nih.gov/17661780/

51. Kurtoglu T, Durmaz S, Akgullu C, Gungor H, Eryilmaz U, Meteoglu I, et al. Ozone preconditioning attenuates contrast-induced nephropathy in rats. Journal of Surgical Research. 2015;195(2):604–11. Available from: https://www.sciencedirect.com/science/article/pii/S0022480415000700

52. Wang L, Chen H, Liu XH, Chen ZY, Weng XD, Qiu T, et al. The protective effect of ozone oxidative preconditioning against hypoxia/reoxygenation injury in rat kidney cells. http://dx.doi.org/103109/0886022X2014950934. 2014;36(9):1449–54. Available from: https://www.tandfonline.com/doi/abs/10.3109/0886022X.2014.950934

53. Meng W, Xu Y, Li D, Zhu E, Deng L, Liu Z, et al. Ozone protects rat heart against ischemia-reperfusion injury: A role for oxidative preconditioning in attenuating mitochondrial injury. Biomedicine & Pharmacotherapy. 2017;88:1090–7. Available from: https://www.sciencedirect.com/science/article/abs/pii/S0753332216320030

54. Orlandin JR, Machado LC, Ambrósio CE, Travagli V. Ozone and its derivatives in veterinary medicine: A careful appraisal. Veterinary and Animal Science. 2021;13:100191.

